# Reliability of Mouse Behavioural Tests of Anxiety: a Systematic Review and Meta-Analysis on the Effects of Anxiolytics

**DOI:** 10.1101/2021.07.28.454267

**Authors:** Marianna Rosso, Robin Wirz, Ariane Vera Loretan, Nicole Alessandra Sutter, Charlène Tatiana Pereira da Cunha, Ivana Jaric, Hanno Würbel, Bernhard Voelkl

**Affiliations:** Division of Animal Welfare, University of Bern, Bern, Switzerland

## Abstract

Animal research on anxiety and anxiety disorders relies on valid animal models of anxiety. However, the validity of widely used rodent behavioural tests of anxiety has repeatedly been questioned, as they often fail to produce consistent results across independent replicate studies using different study populations or different anxiolytic compounds. In this study, we assessed the sensitivity of behavioural tests of anxiety in mice to detect anxiolytic effects of drugs prescribed to treat anxiety in humans. To this end, we conducted a pre-registered systematic review of studies reporting tests of anxiolytic compounds against a control treatment using common behavioural tests of anxiety in mice. PubMed and EMBASE were searched on August 21^st^ 2019 for studies published in English and 814 papers were identified for inclusion. Risk of bias was assessed based on Syrcle’s risk of bias tool and the Camarades study quality checklist on a randomly selected subsample of 180 papers. Meta-analyses on effect sizes of treatments using standardized mean differences (Hedges’ g) showed that only two of 17 test measures reliably detected effects of anxiolytic compounds other than diazepam. Further, we report considerable variation in both direction and size of effects of most anxiolytics on most outcome variables, indicating poor replicability of test results. This was corroborated by high heterogeneity in most test measures. Finally, we found an overall high risk of bias. Our findings indicate a general lack of sensitivity of common behavioural tests of anxiety in mice to anxiolytic compounds and cast serious doubt on both construct and predictive validity of most of those tests. The use of animals to model human conditions can be justified only if the expected results are informative, reproducible, and translatable. In view of scientifically valid and ethically responsible research, we call for a revision of behavioural tests of anxiety in mice and the development of more predictive tests.

## INTRODUCTION

Animal experiments are a key component of basic and preclinical research, where the mechanisms of diseases are studied and new compounds for their treatment are examined for safety and efficacy before being tested in humans (fda.gov). However, the use of animals for research can only be justified when the results obtained are informative (1–3), replicable* (4–6), and translatable* (7,8). Furthermore, public concern for animal welfare urges scientists to comply with the 3Rs principle (9), that is to refine, reduce, or replace the use of animals whenever possible (10,11).

To achieve these goals and ensure responsible scientific practice, the validity* of animal models in use is pivotal (2,12–14). A growing body of evidence indicates the lack of validity of animal models as a potential cause for translational failure (13,15–17). Translational failure can slow down medical advancement in the treatment of human disorders (18–20), put patients in clinical trials at risk (3), waste research resources (21), and harm animals for inconclusive research.

Anxiety disorders are amongst the most common mental health conditions, requiring still new and better treatments (22–26). To study anxiety and to test the efficacy of anxiolytic compounds behavioural tests in mice and other animals are commonly used (22,23,27,28). Such tests are mostly based on exploiting an approach-avoidance conflict, i.e. the conflict an animal may experience between exploring a new, and avoiding a potentially threatening, environment (27,29,30). Amongst the various behavioural tests for rodents, the open-field test is arguably the most popular one (23). This test, although with several modifications (31,32), generally consists of a brightly illuminated arena, enclosed by walls. During the test, an animal is placed inside the arena and behavioural outcomes are recorded. The test was originally established to assess emotionality in rats, using urination and defecation as measures of timidity (31,33). The use of the open-field test was then extended to assess a wider range of behavioural features and psychiatric conditions (27) and adopted for other species. Similar to rats, early studies which employed the open-field test in mice measured defecation and freezing to assess genetic differences in behaviour (34,35). Additionally, the distance travelled in the open-field test has been introduced and--since then--widely used as a measure of locomotor activity to assess, for instance, the effect of sedative or stimulant drugs (36). Further, thigmotaxis in the open-field, namely the tendency to explore the proximity of the walls while avoiding the centre of the arena, is often recorded and interpreted as a proxy for anxiety (27,32,37).

Similar to the open-field test, the elevated plus maze test (38) and the light-dark box test (39) are based on the conflict between the exploration of a new environment and the natural aversion of rodents to bright and open spaces. The rationale behind these tests as measures of anxiety rests on the assumption that a state of anxiety should modulate the animals’ behaviour by reducing exploration, therefore reducing the exposure to (potential) threats (22,27,40). Accordingly, the efficacy of anxiolytic compounds is assessed based on whether and to what extent they attenuate the reduction of exploratory behaviour by the test situation. Other popular tests, such as the hole-board test (41), the elevated zero maze (42), the social interaction test (43), the novelty suppressed feeding test (44), and the four-plate test (45), are based on the same conceptual rationale.

Over the years, behavioural tests for anxiety have been considered validated, because of reported behavioural changes elicited by benzodiazepines, and specifically diazepam (46–48). However, anxiolytic agents such as benzodiazepines also possess anti-depressant and sedative effects, which implies that the observed behavioural effects may not necessarily be due to a change in anxiety, but could be a result of the sedative properties of the drug (36).

Despite their popularity, several experimental studies, as well as literature reviews, have highlighted inconsistent results in the behavioural outcomes elicited by new classes of anxiolytics, therefore questioning the suitability of these outcomes as indicators for anxiety (29,36,46,49,50). Benzodiazepines, although popular in the past to treat anxiety, have now been replaced by better pharmacological compounds with fewer side effects and lower withdrawal-related risks (51–53). Selective Serotonin Reuptake Inhibitors (SSRIs) or Serotonin–Norepinephrine Reuptake Inhibitors (SNRIs), which are now used as a first-line pharmacological treatment for human anxiety disorders, have failed to give reliable results in rodent behavioural tests of anxiety (29,36,46,50,54).

Here, we aimed to assess the validity of common behavioural tests of anxiety in mice by evaluating their responsiveness to anxiolytic compounds prescribed to humans, a process known as ‘reverse translation’ (55,56). To this end, we performed a pre-registered systematic review of research papers that had used these tests on laboratory mice, for a broad range of anxiolytic compounds. We investigated the overall effect size for a range of test measures of common behavioural tests as well as the variation of the reported outcomes across the published literature. Additionally, we evaluated sample heterogeneity and estimated the quality of reporting through a risk of bias assessment.

**\*Glossary of key terms**

1. **Replicability**: the likelihood with which results can be replicated by an independent study.
  - Relevant literature: (5,6,57–61)
2. **Translatability**: the extent to which results obtained in an animal model can be replicated in the system which is being modelled.
  - Relevant literature: (16–18,62–64)
3. **Validity**: to be fit for use in research, and therefore be considered to be a valid animal model, a test or animal model should meet several criteria of validity, including:
  i. **Construct validity**: the extent to which the test can measure what it is supposed to measure
  ii. **Predictive validity**: the extent to which a test can predict a certain outcome in the system that is being modelled.
  - Relevant literature: (1,2,12,28,65,66)

## METHODS

### PRE-REGISTRATION

Prior to data extraction, in November 2019, this study was pre-registered at SYRCLE (see supplementary information for the pre-registration protocol).

### SEARCH STRATEGY

The search strategy consisted of i) a list of anxiolytic compounds, ii) the keyword “mice”, and iii) a list of behavioural tests for anxiety. To define the list of anxiolytic compounds, we used a combination of the following databases to list compounds that are commonly used to treat anxiety disorders in humans: DrugBank (drugbank.ca); FDA Drug Approval Databases (fda.gov); Anxiety and Depression American Association (adaa.org). We selected the following compounds: alprazolam, amitriptyline, buspirone, chlordiazepoxide, citalopram, clomipramine, clonazepam, clorazepate, desipramine, diazepam, doxepin, duloxetine, escitalopram, fluoxetine, flurazepam, fluvoxamine, hydroxyzine, imipramine, lorazepam, maprotiline, mirtazapine, nortriptyline, oxazepam, paroxetine, protriptyline, sertraline, temazepam, trazodone, triazolam, trimipramine, venlafaxine. A literature search allowed us to identify behavioural tests commonly used to assess anxiety in mice (Table 1). Each test that yielded more than 10 results, when searched on PubMed (on date July 15^th^ 2019) in combination with the aforementioned list of compounds, and the keyword “mice”, was included in the search (Supplement 1). The search was performed on PubMed (ncbi.nlm.nih.gov/pubmed) and EMBASE (embase.com), on August 21st, 2019.

**Table 1.**
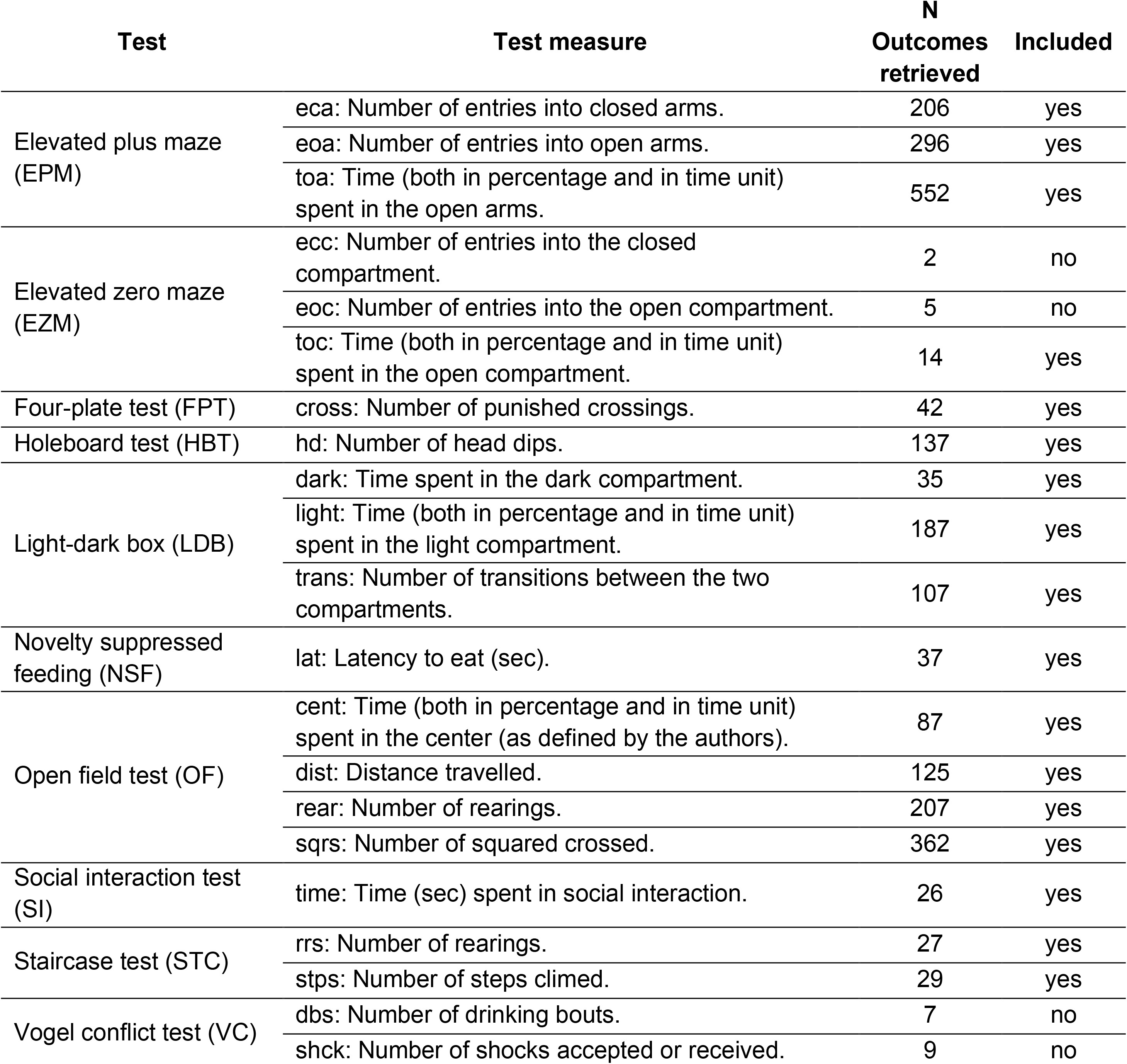
Behavioural tests for anxiety in mice and relative test measures included in the search.

### STUDY SELECTION

After reference retrieval, we excluded paper duplicates using the reference manager software Citavi 6.4 (Swiss Academic Software GmbH, Wädenswil, CH). The main reviewer (MR) scanned the titles, abstracts and/or methods of these papers, and excluded all those, which did not use the behavioural tests of interest (Table 1), mice, or the selected anxiolytic compounds. Additionally, we excluded papers that were not original research papers and papers that were not written in English. After the first scan, two independent reviewers (main reviewer: MR, second reviewers: RW, AL, NS) performed the full paper screening and the data extraction.

### STUDY CHARACTERISTICS

Studies were included or excluded according to the pre-specified inclusion/exclusion criteria (Supplement 1). For each paper, two reviewers independently extracted information about the animals (i. strain, ii. sex, iii. age, iv. transgenic ID; v. stress or defeat treatment), about the treatment (vi. compound, vii. dosage, viii. route of administration, ix. time of administration before testing), and about testing (x. open field size, xi. test duration). For each test, we selected test measures suggested by the authors as measures of anxiety (Table 1). For each test measure, we extracted mean values, sample size, and either standard deviation or standard error of the mean, for both treatment and control group. We accepted any control group as declared by the authors (e.g. administrating water, saline solution, etc.). Information from graphical data was extracted using the online software Automeris (https://apps.automeris.io/wpd/).

### DATA ANALYSIS

The statistical analysis was performed in R (1.4.1103) (67) with the package metafor 2.4-0 (68). For each study, we computed the standardized mean difference Hedges’ g between the control and the treatment group as the chosen indication of effect size (metafor::escalc). We included any test measure that yielded at least 10 results. Consequently, four measures (EZM-*eoc*, EZM-*ecc*, VT-*shcks*, VT-*dbs*) were excluded from further analysis. For the measures LDB-*dark*, EPM-*eca*, NSF-*lat*, STC-*rrs* we reversed the sign of the effect size, because a decrease in behaviour manifestation is expected as a result of treatment. Our data pool was subset by test measure and a meta-regression model was fitted for every subset.

rma (yi, vi, mods= ~ factor (compound) - 1, random = list(~ 1 | study/observation,~ 1 | strain) Standardized mean differences (Hedges’ g) were tested with the modifier ‘compound’ (anxiolytic compounds) against the null hypothesis of the estimated effect size for each compound group equalling zero. Publication and strain were added as random effects. To assess the overall estimated effect size, independent of anxiolytic compound, the same model syntax was used, excluding the factor modifier. Total and partial I^2^, indicating the percentage of sample variation, were used as a measure of heterogeneity, and were calculated using the methods proposed in (69).

### RISK OF BIAS

Due to the large sample size, an assessment of quality was made on a subsample consisting of 180 randomly selected papers. The assessment was done by two independent reviewers (MR, CP), who evaluated 80 different papers each, as well as 20 papers that were reviewed by both investigators, to estimate inter-rater reliability. We used an adapted combination of the CAMARADES study quality checklist and SYRCLE’s risk of bias tool (Supplement 1).

## RESULTS

### STUDY SELECTION

Our search retrieved 744 papers from PubMed and 2533 papers from EMBASE of which 1764 were excluded in the first steps of the review (Fig 1). In particular, 533 were excluded as paper duplicates, and 1231 were excluded based on abstract and/or method section screening. The full texts of 1513 papers were screened and 814 of those papers were included in the data extraction process according to the pre-specified criteria. As the search strategy identified key words in all fields of the text, several papers not relevant to us were identified; 331 papers were excluded because the sample size was unclear or not reported, 62 papers were excluded because the text was unavailable publicly, 59 papers were excluded because compounds other than the ones of interest were used, or compounds were used in combination with other compounds, 48 papers were excluded because of issues in the reporting of the outcomes, 40 papers were excluded because they had formats other than research papers, 33 papers were excluded because the behavioural tests used were different from the ones of interest, 25 papers were excluded due to ambiguity regarding the measure of variance of the reported outcomes, 24 papers were excluded because they used animals other than mice, or because of ambiguity in the species of animal used, and 13 papers were excluded for other reasons (i.e. missing controls, treatment administered to mothers, etc.).

**Fig 1.**
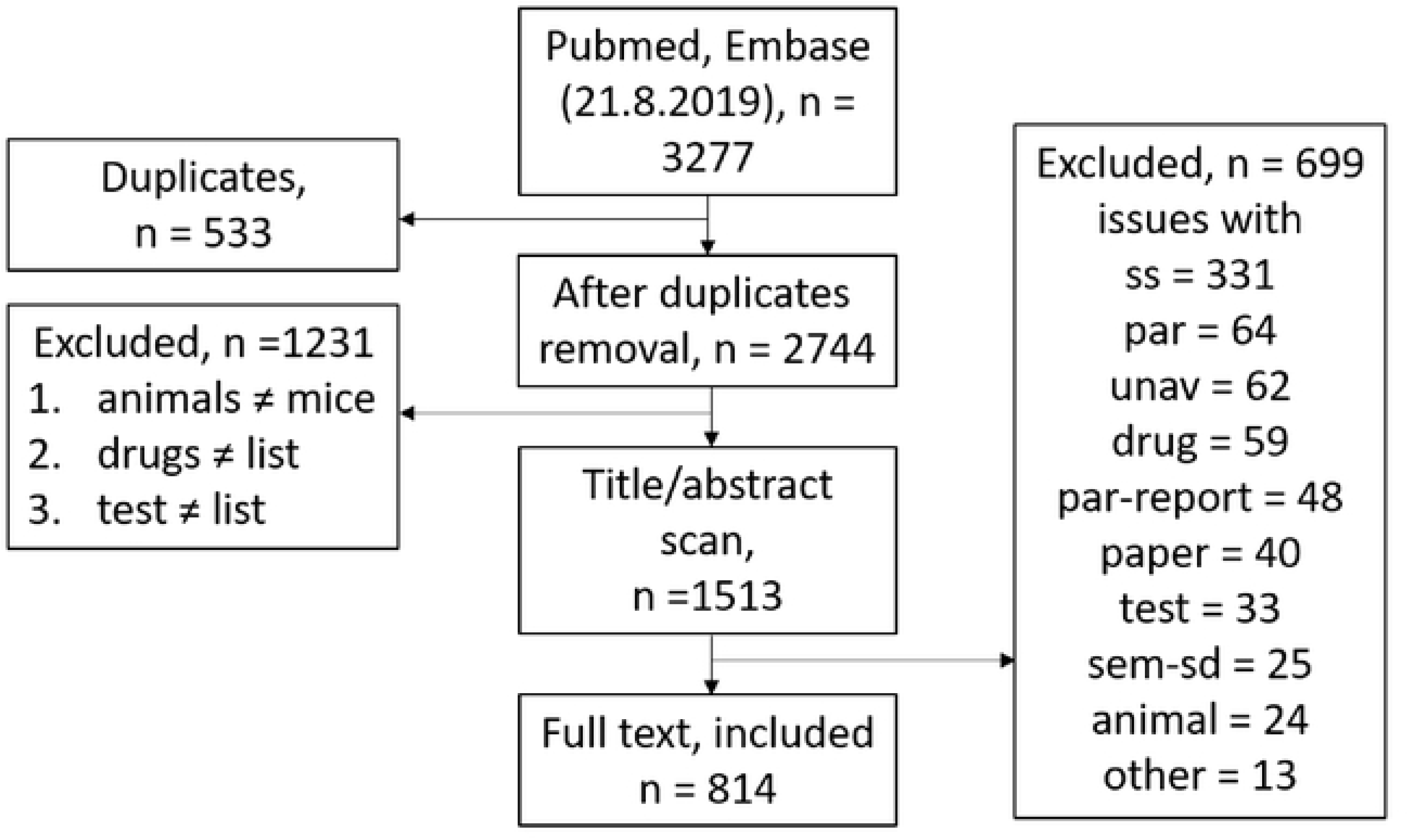
Flowchart of the screened papers and reasons for exclusion. ss: unclear or absent sample size; unav: paper unavailable; par: incompatible outcomes reported; drug: incompatible compounds used; par-report: issues with the reporting of the outcomes; paper: wrong format of paper; test: incompatible behavioural test used; animal: wrong animals used; sem-sd: unclear or absent measure of variance; other.

### STUDY CHARACTERISTICS

All the eligible studies used mice, which were tested in behavioural tests after administration of anxiolytic compounds. The Supplementary table illustrates the details of data distribution in the different test measures of interest in combination with each compound. Due to reporting of multiple outcomes per paper, a total of 2476 outcomes were distributed across 17 different test measures, in combination with 25 different anxiolytic compounds. The test measures from the elevated plus maze and the open field made up the great majority of outcomes (74%, Table 1), followed by the light-dark box test and the holeboard test contributing a total of 13% and 5% of the outcomes, respectively. A minor contribution was attributed by the staircase test (the staircase test, n = 56, “rrs” n = 27, “stps” n = 29), the four-plate test (n = 42), the novelty suppressed feeding test (n = 37), the social interaction test (n = 26), and the elevated zero maze (n = 14). The great majority of these measures were recorded when used in combination with benzodiazepines (72%), with diazepam being the most frequently used compound (65%). SSRIs was the second most common compound class (20%), with fluoxetine (12%) being its most frequently used representative.

### RISK OF BIAS

A sub-sample of 180 papers was analysed in detail to assess the risk of bias across 17 different items (Table 2). All the scored papers were published in peer-reviewed journals, and most of them reported mouse strain (95%), sex (90%) and housing temperature (75%). 31% of the papers reported details regarding compliance with animal welfare regulations, 43% of the papers reported details on the statistical analysis, and 34% of the papers reported details on the blinding procedures. For the following five items, we scored a high risk of bias: automatic allocation to treatment group (97%), randomized order of testing (92%), a-priori sample size calculation (98%), random housing (95%), and blinding of investigators (95%). Further details are reported in Table 2.

**Table 2:**
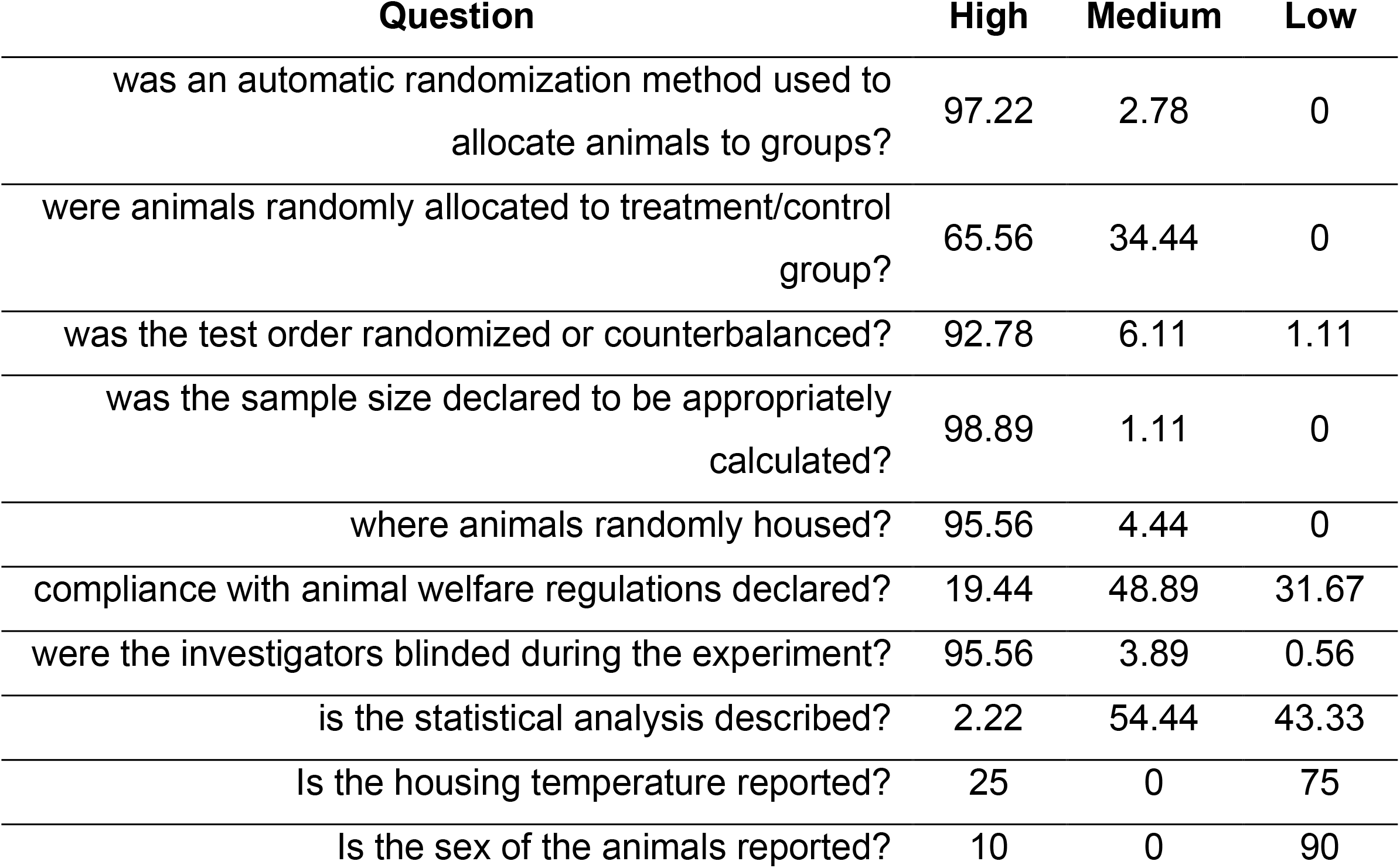
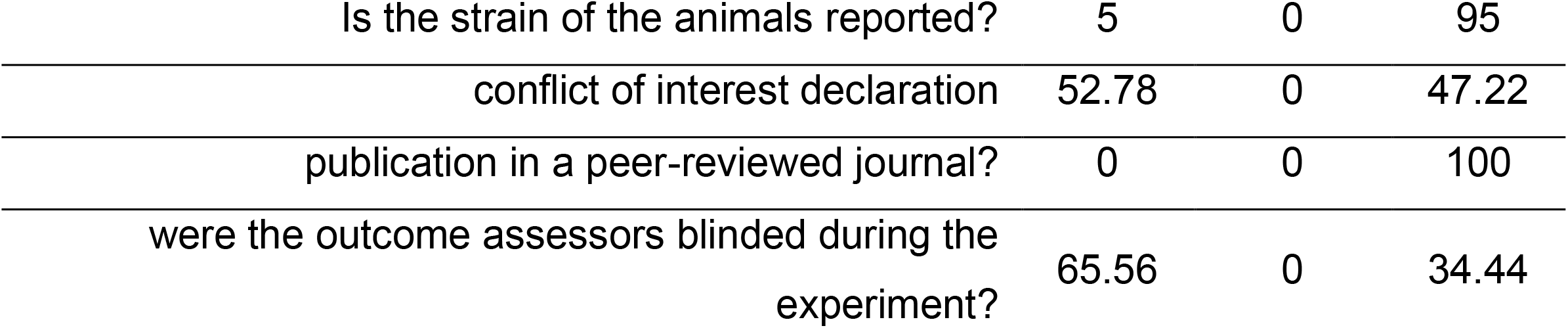
Results of the risk of bias assessment. Values in the table indicate percentages of papers, which scored either as high, medium, or low risk of bias in each item (row).

### SYNTHESIS OF RESULTS

Estimated effect sizes varied greatly across the majority of the test measures and compounds (Fig 2). The overall estimated effect size allows determining whether there is evidence of an anxiolytic effect on the behavioural measures elicited by a range of anxiolytic compounds. Ten out of the 17 test measures yielded a positive overall effect size significantly different from zero (EPM-*eca*, EPM-*eoa*, EPM-*toa*, FPT-*cross*, LDB-*light*, LDB-*trans*, NSF-*lat*, OF-*cent*, SI-*time*, STC-*rrs*), while overall effects of the remaining seven did not significantly deviate from zero.

**Fig 2:**
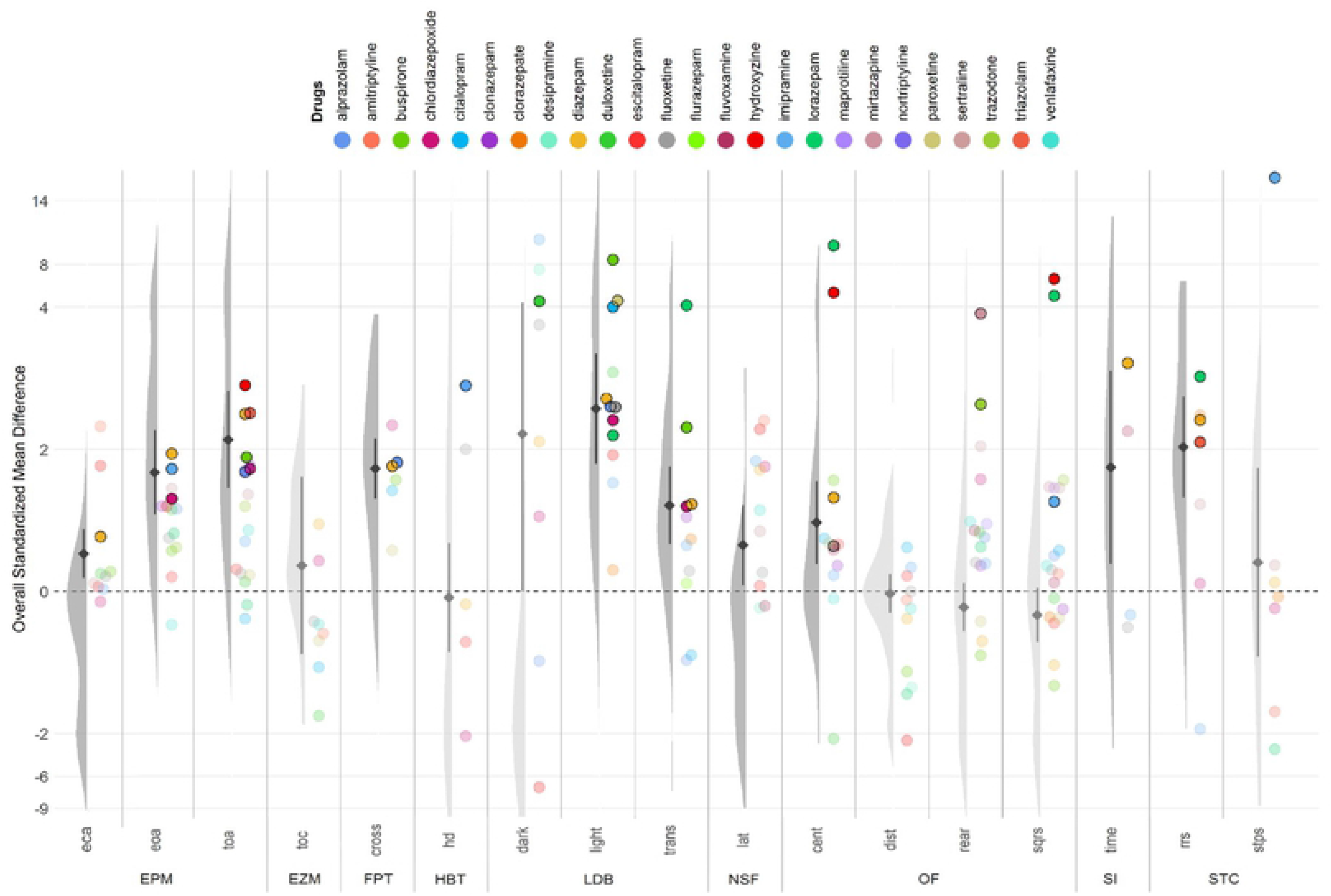
Violin plots showing the probability density distribution of the calculated effect size (x-axis) of the individual studies for each test measure. Overlapped to the violin plots, the overall estimated effect size for each test measure, indicated by the diamonds, and the relative 95% confidence interval. Points indicate the estimated mean effect size for each compound. Colours indicate anxiolytic compounds. Opacity is applied to not significant effect sizes, i.e. the lower bound of the 95% confidence interval is lower than zero. An interactive version of the Fig can be found online at https://mrossovetsuisse.shinyapps.io/Shiny_SR/.

For each meta-analysis, the factor ‘compound’ was tested for significance to assess whether any of the anxiolytic compounds affected behavioural outcomes. For this, the null hypothesis to be tested assumes the estimated effect sizes for all compounds to be zero (68). After family-wise correction for multiple testing for the 17 meta-analyses performed, five measures showed no significant effect, namely EZM-*toc*, LDB-*dark*, NSF-*lat*, OF-*dist*, and SI-*time* (Table 3).

**Table 3:**
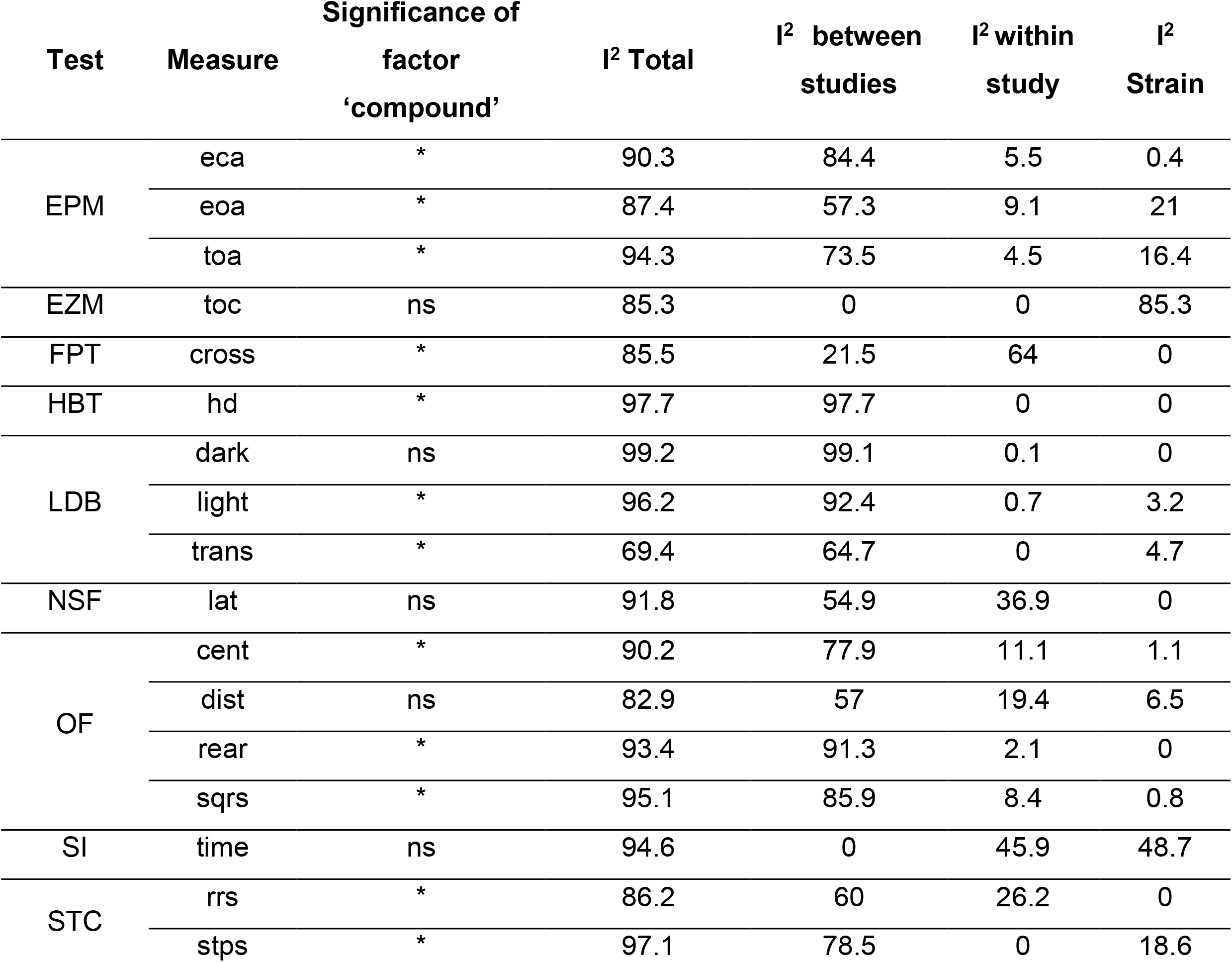
Significance level of moderator effect (treatment × compounds interaction), total and partial I^2^ estimates per test measure.

| Test | Measure | Significance of factor 'compound' | I<sup>2</sup> Total | I<sup>2</sup> between studies | I<sup>2</sup> within study | I<sup>2</sup> Strain |
| --- | --- | --- | --- | --- | --- | --- |
| EPM | eca | * | 90.3 | 84.4 | 5.5 | 0.4 |
|  | eoA | * | 87.4 | 57.3 | 9.1 | 21 |
|  | toa | * | 94.3 | 73.5 | 4.5 | 16.4 |
| EZM | toc | ns | 85.3 | 0 | 0 | 85.3 |
| FPT | cross | * | 85.5 | 21.5 | 64 | 0 |
| HBT | hd | * | 97.7 | 97.7 | 0 | 0 |
| LDB | dark | ns | 99.2 | 99.1 | 0.1 | 0 |
|  | light | * | 96.2 | 92.4 | 0.7 | 3.2 |
|  | trans | * | 69.4 | 64.7 | 0 | 4.7 |
| NSF | lat | ns | 91.8 | 54.9 | 36.9 | 0 |
| OF | cent | * | 90.2 | 77.9 | 11.1 | 1.1 |
|  | dist | ns | 82.9 | 57 | 19.4 | 6.5 |
|  | rear | * | 93.4 | 91.3 | 2.1 | 0 |
|  | sqrs | * | 95.1 | 85.9 | 8.4 | 0.8 |
| SI | time | ns | 94.6 | 0 | 45.9 | 48.7 |
| STC | rrs | * | 86.2 | 60 | 26.2 | 0 |
|  | stps | * | 97.1 | 78.5 | 0 | 18.6 |

For each test measure, we calculated total and partial I^2^ as a measure of heterogeneity. For 15 out of 17 measures, total I^2^ was above 85%. The partial I^2^ attributed to ‘strain’ contributed little to the total I^2^, except for SI-*time*, where it accounted for 48% of the total heterogeneity. Partial I^2^ attributable to within-study heterogeneity varied greatly across measures: in 10 cases being <10%, while being more pronounced in others (e.g. 64% for FPT-*cross*). Between-study heterogeneity explained the greater part of the total heterogeneity for 14 out of the 17 measures (Table 3).

Given the 25 compounds and 17 test measures, there are a total of 425 compound-by-measure combinations. We found reported study outcomes for 182 of those compound-by-measure combinations (details summarized in the Supplementary Table). The number of outcomes per combination varied from 1 to 413, with 118 compound-measure combinations with more than one outcome recorded. Of these, only 32 had a positive and significant effect size (i.e. the lower bound of the 95% confidence interval being larger than zero), while 86 combinations did not show a positive effect (Fig 2 and Supplementary Table). Diazepam was the compound that elicited a significant positive effect size in 9 out of 17 test measures. Overall, most of the combinations with a significant effect size were due to benzodiazepines, with 20 positive effects out of 32. LDB-*light* yielded a positive effect size for most of the anxiolytic compounds tested, 8 out of 11, and EPM-*toa* yielded a positive effect size for 5 out of 15 anxiolytic compounds. The rest of the test measures detected an effect for at most two anxiolytic compounds, across the range with which they were tested.

The percentage of individual observations that detected a positive significant effect varied greatly across the different combinations of test measures and anxiolytic compounds, ranging from 0% to 100% (Table 4). As all the compounds included in this analysis have been shown to reduce anxiety in humans, we assessed the sensitivity of behavioural tests outcomes to detect the expected anxiolytic effect of these compounds in mice based on the logic of reverse translation. Thus, we used the proportion of individual studies reporting a significant positive effect as a measure of sensitivity and an estimate of the true positive rate. To conclude that a behavioural test reliably detects an anxiolytic effect, we require that individual studies detect significant effects (positive effect size with a 95% confidence interval not including zero) in at least three out of four cases (i.e. 75%). The majority of behavioural measures failed to reliability detect an effect for the majority of the compounds. In 89 out of 118 combinations for which more than one outcome was recorded, less than 75% of individual studies reported significant positive effects, while only for 29 combinations, the proportion was greater than 75%. Table 4 suggests that diazepam was the compound that most often elicited a behavioural change detectable in five test measures. Here, we also observe a higher number of studies as compared to other compounds. Out of the 29 ‘reliable’ combinations, benzodiazepines were the dominant compound class, showing reliable results in 14 combinations. LDB-*light* seems to be the most promising candidate to detect an anxiolytic effect, with the majority of individual studies detecting an effect in seven out of 11 anxiolytic compounds across compound classes. Furthermore, EPM-*eoa* and EPM-*toa* reliably detected effects for 3 and 4 anxiolytic compounds, respectively. Similarly, OF-*sqrs*, reliably detected an effect of 3 anxiolytic compounds, but the number of individual studies was far lower than for the EPM. Forest plots (Fig 3 and Supplementary Material) show how for some measures the estimated effect sizes for individual studies range from highly negative values to highly positive ones, spreading in an almost symmetrical fashion across the null. Clear examples of such pattern can be seen in the forest plots of EPM-*eca*, HBT-*hd*, LDB-*trans*, NSF-*lat*, OF-*dist*, OF-*rear*, OF-*sqrs*, and STC-*stps*.

**Table 4:**
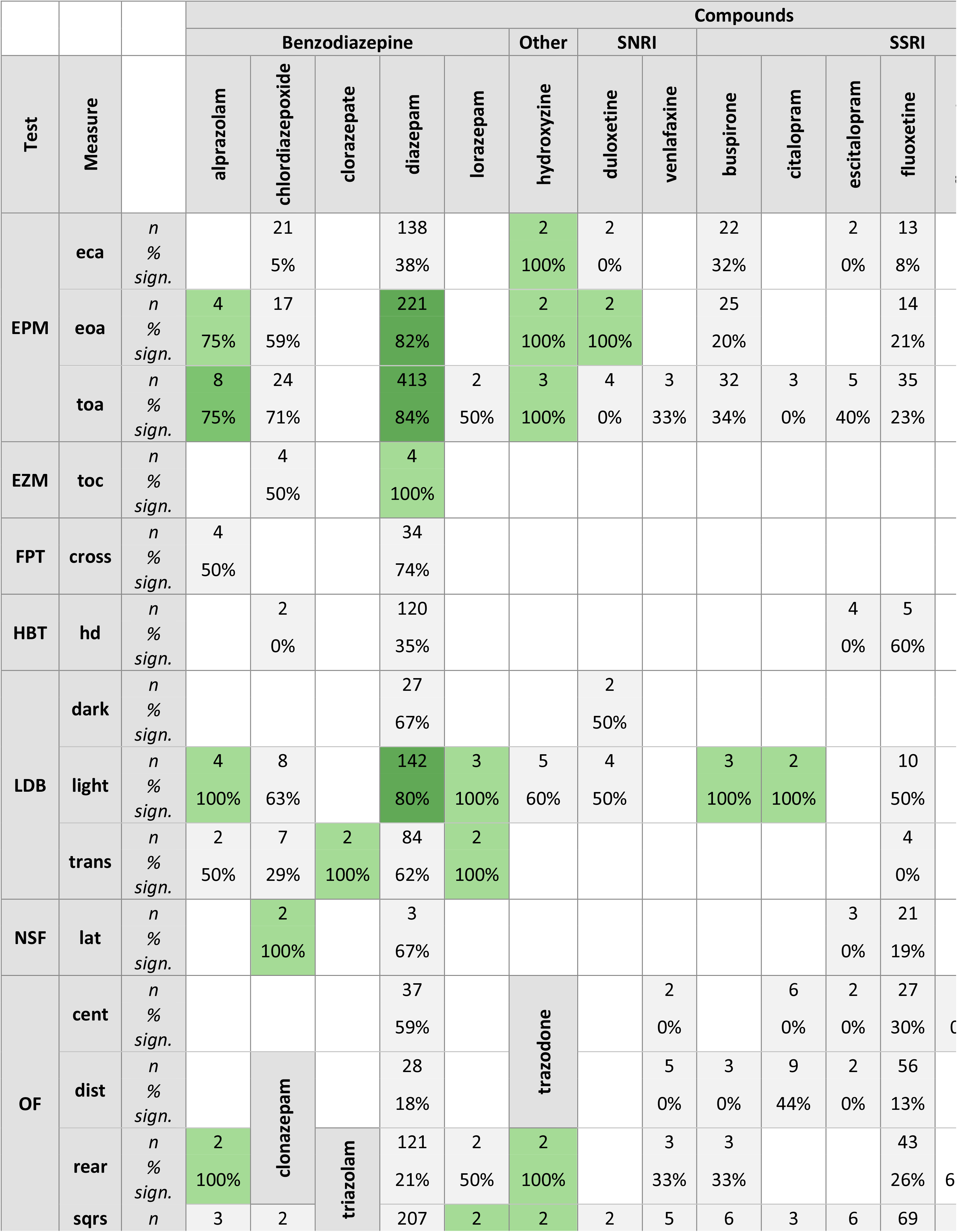
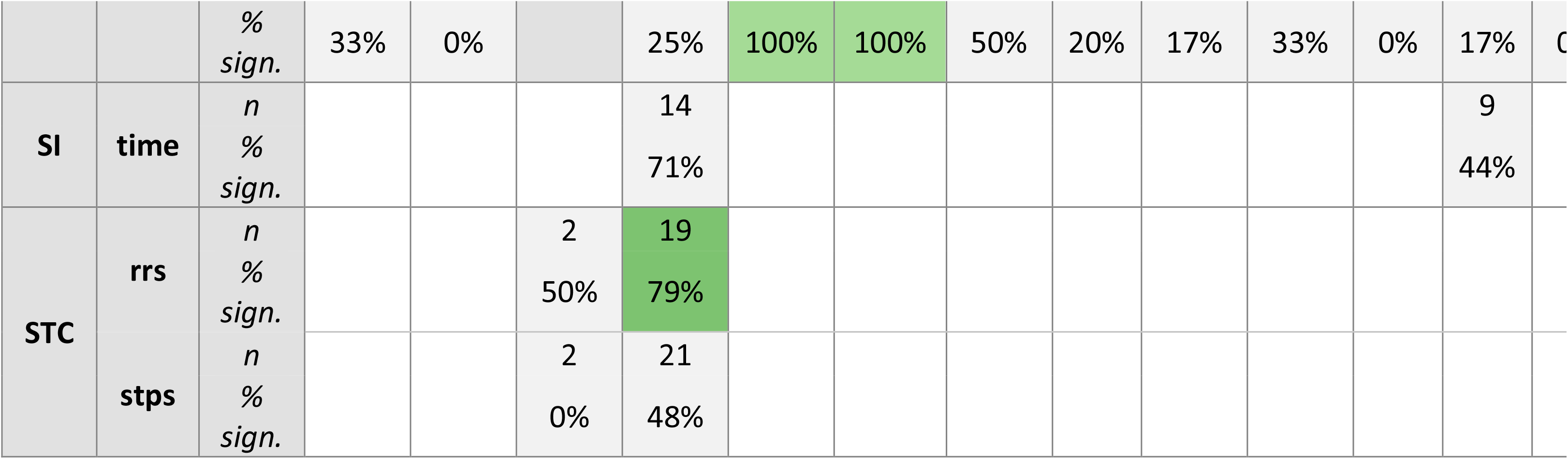
Number of studies and percentage of positive studies, per combination of test measure and anxiolytic compounds. Cells in grey indicate a percentage of positive studies <75%. Coloured cells highlight a percentage of positive studies >75%. Colour gradient indicates an increasing number of studies. Combinations with only one study were excluded from the table.

**Fig 3:**
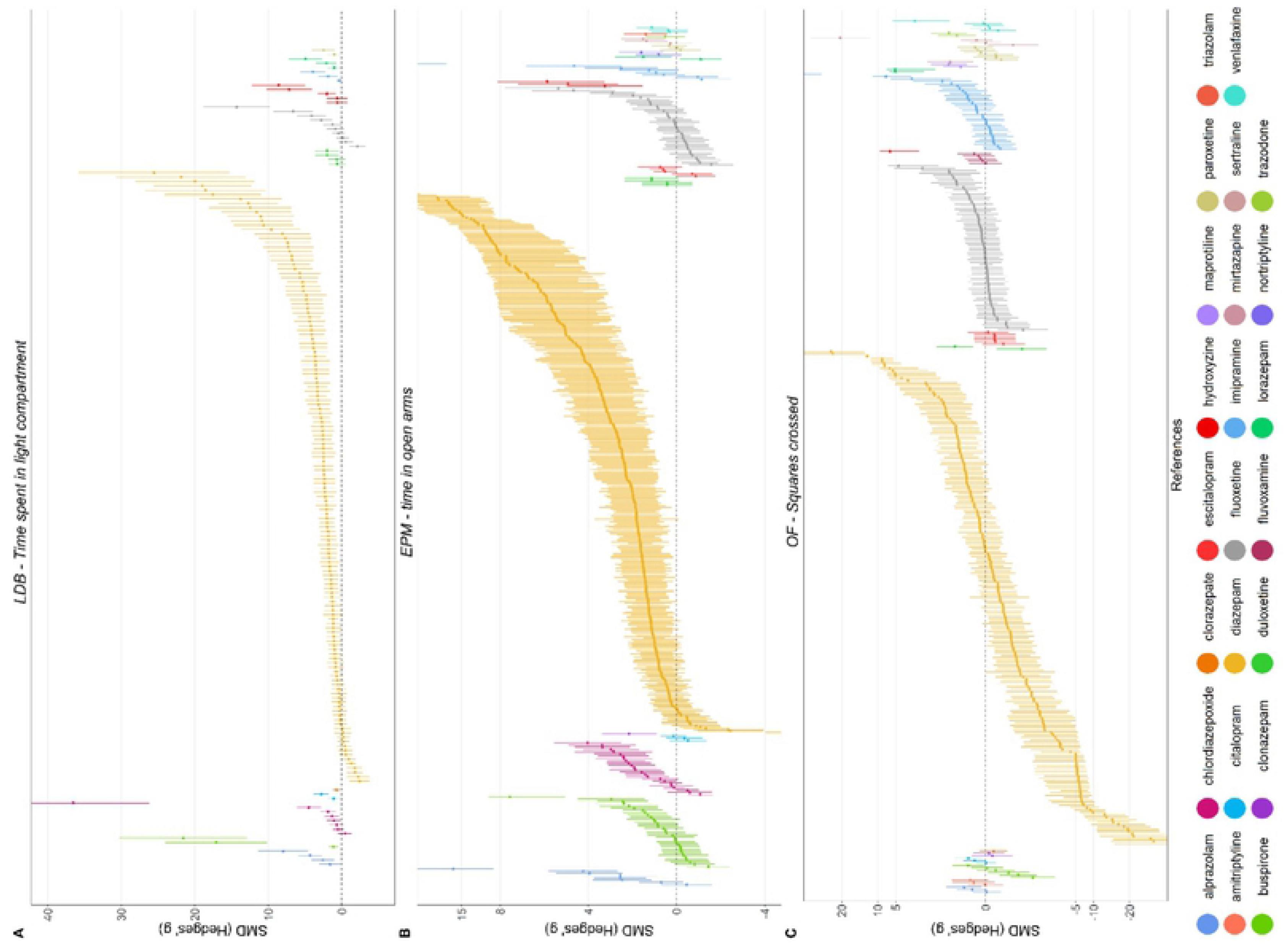
Forest plots of three selected test measures. : A: LDB-*light*, B: EPM-*toa*, C: OF-*sqrs*, sorted for increasing effect size. Different colours indicate different anxiolytic compounds, as indicated in the legend (See Supplementary material for remaining measures).

## DISCUSSION

With the present study, we aimed at providing a synthesis of the reliability of mouse behavioural tests of anxiety. We assessed their sensitivity to a broad range of anxiolytic compounds approved for the treatment of anxiety in humans, using a systematic and unbiased approach. Briefly, we found reported effects to vary greatly across studies and test measures, in addition to overall high heterogeneity and important risks of reporting bias.

We found that for five of the 17 test measures, none of the anxiolytic compounds had a significant effect, whereas, for the remaining 12 test measures, an effect of at least one anxiolytic compound was detected. Additionally, we investigated the overall estimated effect size for each test measure, irrespective of anxiolytic used, and found null or negative overall effects for seven test measures.

For the majority of the test measures and specific compounds, we have observed great variation in the estimated effect sizes, ranging from highly negative to highly positive values, and resulting in estimated cumulative effect sizes close to zero (e.g. in OF-*sqrs* and in OF-*rear*, and in HBT-*hd*.). Additionally, we observed that the effect size estimates of individual studies, which reported a significant effect of a compound also varied greatly even for those combinations in which the overall estimated effect size was positive. Because all of the compounds included in our study were shown to have anxiolytic effects in humans, we consider the proportion of individual studies as a measure of how reliably such behavioural tests can detect behavioural changes elicited by anxiolytic compounds. Overall, only 1254 out of all 2476 contrasts (i.e. 50%) showed significant treatment effects.

Investigation of the total and partial heterogeneity showed that the greater portion (median 74%) of the sample heterogeneity, across test measures, is produced by differences between studies. Such a high level of between-study heterogeneity seems to be common in several fields of animal research (70–73).

There were only two test measures in which the between-study heterogeneity was as low as expected due to random variation alone: SI-*time* and EZM-*toc* These test measures were, however, not sensitive to effects of anxiolytic compounds. Within-study heterogeneity varied greatly across measures but was overall lower than other partial heterogeneity measures, hinting at high levels of standardization within laboratories.

Even though our results show that most of the test measures do not reliably detect behavioural changes elicited by several anxiolytic compounds, we have found two test measures - EPM-*toa* and LDB-*light* - that appear to be sensitive both in terms of detecting a positive effect of anxiolytic compounds and to reliably detect a positive effect in the majority of the individual studies. Additionally, these test measures show significant positive effect sizes for a wider range of anxiolytic compounds than the other measures. With 73% (EPM-*toa*) and 78% (LDB-*light*), respectively, of individual studies reporting a positive effect, the false-negative rates approach the minimally recommended threshold of 0.2. Thus, these measures seem to be promising starting points for refinement and the development of reliable test procedures.

The substantial variation observed between studies using the same test measure and anxiolytic compound with comparable dosages is likely to be attributed to environmental, genetic, and procedural differences. Previous analyses of behavioural test outcomes for the effect of mouse strain on both basal levels of performance and performance after the administration of anxiolytic compounds highlighted substantial strain differences and often conflicting results (46,74–77). Surprisingly, we found only weak effects of mouse strain on heterogeneity for most of the test measures. Apart from genetic background, differences in sex, age, housing conditions, and test environment may contribute to between-study variation. Unfortunately, these are only sporadically and scantily reported. We invite the readers to explore our publicly available dataset through our online application, available at https://mrossovetsuisse.shinyapps.io/Shiny_SR/, which allows displaying data subset by sex, strain, stress treatment and dosage.

Taken together, our results show that most behavioural test measures are unreliable in detecting behavioural changes elicited by anxiolytic compounds other than benzodiazepines and in particular diazepam. This corroborates the previously voiced suspicion that most popular behavioural tests of anxiety are in fact "benzodiazepines tests" (29,47). The behavioural effects elicited by benzodiazepines in these tests have been proposed to reflect disruption of normal behaviour, possibly resulting in altered impulse control rather than attenuated anxiety (47,78).

The behavioural tests included in our study heavily rely on changes in exploration patterns to determine anxiety levels and such test procedures may not be able to disentangle behavioural changes in exploration and anxiety (37,49,79). A clear example of this problem is the open field test, which is sometimes performed to assess anxiety but sometimes to control for locomotor activity in combination with other tests of anxiety (80,81). For example, if the response of animals to a compound is tested in both the LDB and the open field, an increase in LDB-light in the absence of a change in locomotor activity in the open field would suggest that the investigated compound has a specific anxiolytic effect, but no sedative effect, which is highly desirable in anxiolytics especially from a translational perspective (82–84). Upon literature review, we have found as many records in which the open field was performed as a test of locomotor activity (80,81,85,86), as we have found records in which it was performed as a test of anxiety (87–90). Here, we identify an issue with the continuation of such tests as long-held standards that may not be appropriate, due to the researcher’s degree of freedom in the interpretation of the test’s meaning (91,92).

On a different note, our findings question the standard classification of effect sizes in animal behavioural research. Cohen introduced what are, up to date, considered the conventional thresholds for small, medium, or large effect sizes (namely, a Cohen’s d of 0.2, 0.5, and 0.8 respectively (93)). The author warned for caution (p. 25) in using these thresholds for power analysis outside the scope of the field for which they were initially thought for (psychology or sociology). Study populations of laboratory animals are normally characterised by high degrees of both genetic and environmental standardization (94–96). Therefore, populations of animal studies are usually much more homogenous, producing much lower levels of random variation, when compared to study populations of clinical studies (97). This difference has important implications for the interpretation of standardized effect sizes like Cohen’s d or Hedges’ g. Due to the higher level of standardization in animal studies and the resulting low within-group variation, a given mean difference between a control and a treatment group will result in a much higher standardized effect size. For example, for EPM-*toa,* (98) reported 123.8 seconds spent in the open arms for the control group and 207.3 seconds for the group receiving diazepam. Given the corresponding standard errors of 0.4 and 0.7 for the control and the treatment group, respectively, this amounts to a standardized effect size of 40.6, which is on an entirely different scale of magnitude than a Cohen’s d of 0.8, the reference for “large” effects. While this is one of the more extreme examples, we note that EPM-*toa* had an average effect size across drugs of 2.13, with 77% of the total studies reporting an effect size larger than the standard large effect of 0.8. Correct estimation of expected effect sizes is essential for proper power analyses and sample size calculations, with important implications for animal welfare. Considering the large achieved effect sizes, the power analyses based on the “standard Cohen’s values” are likely to lead to unnecessarily large required sample sizes. Because of this, we call for a cautious interpretation and more contextualized use of effect size classification, according to each field of research.

Our risk of bias assessment showed overall high-risk scores for most of the items. Although the common checklists and tools for risk of bias analyses assess reporting quality rather than study quality, high risks of bias can have serious implications for the reproducibility and replicability of study findings. Albeit efforts have been made to develop more stringent guidelines for both designing and reporting of animal studies (99,100), we observed an overall low quality of reporting, which likely reflects poor study design and conduct. For instance, researchers failed to report the sex or the strain of the animals in 10% of the cases, and important aspects of the housing conditions (e.g. light intensity and temperature), randomization and blinding procedures, testing conditions (e.g. apparatus size, light intensity, and time of testing), as well as sample size calculations were reported only sporadically.

Our study re-evaluates the suitability of behavioural tests of anxiety in mice, showing low to no sensitivity to anxiolytic compounds (other than diazepam) commonly used for the treatment of anxiety in humans. These finding let us expect poor predictive validity for the discovery of new compounds to treat anxiety disorders in humans and points at a high false-negative rate for individual studies. Additionally, our results highlight considerable idiosyncrasy in the results of the behavioural tests as they are currently performed, with the majority of the tests producing irreproducible and often contradicting results. These findings are corroborated by previous evidence for poor replicability of behavioural tests for anxiety (46,47). Animal tests that lack replicability and validity do not generate new knowledge and, consequently, lose their ethical justification. Additionally, invalid pre-clinical animal trials impair scientific and medical advancement, impacting human subjects in need of treatment. Following the 3Rs principle, effort must be made to improve the quality of animal models for anxiety by developing more informative and reproducible tests with a sound rationale producing results of high internal as well as external validity. This can lead not only to a significant improvement of experimental results but also to more comprehensive and conclusive evidence synthesis in systematic reviews, tackling the prominent bias for positive publications.

## Acknowledgements

The authors would like to thank Dr. Cathaljin Leenaars and Dr. Georgia Salanti for their assistance in the data analysis and their valuable feedback on the interpretation of the results.

